# Quantification of X-chromosome inactivation in fibroblast and iPSC models of *UBQLN2* ALS/FTD using allele-selective qPCR

**DOI:** 10.64898/2026.08.14.744950

**Authors:** David C. Gordon, Kyrah M. Thumbadoo, Serey Naidoo, Agnes L. Nishimura, Miriam Rodrigues, Harry Fraser, Anthony N. Cutrupi, Richard H. Roxburgh, Christopher E. Shaw, Marina L. Kennerson, Emma L. Scotter

## Abstract

Pathogenic missense variants in the X chromosome gene *UBQLN2* cause amyotrophic lateral sclerosis (ALS), often accompanied by frontotemporal dementia (FTD). As an X-linked gene, *UBQLN2* is subject to X chromosome inactivation (XCI), a process wherein one X chromosome in each cell is randomly inactivated to a Barr body throughout the body in females, creating a mosaic of allelic expression in the tissues of heterozygotes. Despite heterozygous females constituting a majority of reported cases of *UBQLN2-*linked ALS/FTD, and the known influence of XCI on neurological disorders at large, no current disease models account for XCI.

Here we report the characterisation of 12 iPSC clones carrying the ALS/FTD-causing p.T487I (c.1460C>T) *UBQLN2* variant. These clones, originally derived from 3 heterozygous carrier fibroblast lines, underwent validation of homeostatic Barr body retention. Erosion of XCI in a subset of the lines was correlated with biallelic expression (of both wildtype and mutant *UBQLN2*), as measured through a novel allele-selective qPCR (AS-qPCR) assay and verified by amplicon-based Illumina sequencing and Sanger chromatogram quantification, enabling selection of iPSC clones best retaining XCI. Together, this *UBQLN2* AS-qPCR assay and selected iPSC clones will enable studies of the role of XCI and its skew in female resilience to *UBQLN2* p.T487I-linked ALS/FTD and enable development of allele-selective therapies.

## Introduction

Pathogenic missense variants in the X chromosome gene *UBQLN2* cause amyotrophic lateral sclerosis (ALS), often accompanied by frontotemporal dementia (FTD) (1). Variants in *UBQLN2* are the only known X-linked dominant causes of ALS. As such, *UBQLN2*-linked ALS /FTD demonstrates the hallmarks of diseases associated with X chromosome inactivation (XCI) in females (2). Such hallmarks include, on average, older age of onset and death and lower neuropathological burden in females, but with high variability (3, 4). Members of an Australasian pedigree carrying a pathogenic *UBQLN2* variant c.1460C>T (p.T487I) (5) demonstrate this signature of female resilience to X-linked ALS/FTD (2). In contrast, and supporting a role for XCI, these hallmarks are not recapitulated by transgenic animal models that overexpress *UBQLN2* independent of the genomic context of the X chromosome (6-21). Therefore, fibroblasts and induced pluripotent stem cells (iPSCs) derived from *UBQLN2* variant c.1460C>T (p.T487I) donors, combined with accurate methods to quantify expression of each *UBQLN2* allele, should enable better modelling of the role of XCI as a modifiable determinant of disease.

X chromosome inactivation occurs in early embryogenesis in human females, enabling dosage compensation between cells through stochastically silencing either the maternal or paternal X chromosome (22). The inactivated X chromosome is silenced through the exclusion of activators and enrichment suppressors of gene expression, on both the genomic DNA and chromatin, across the silenced X chromosome. One canonical marker of such repression is trimethylation of lysine 27 of histone 3 (H3K27me3) (23). Enrichment of repressive markers is associated with formation of a single compact and largely transcriptomically inert ‘Barr body’ (24). All daughter cells inherit this randomised epigenetic footprint to produce a mosaic of maternal and paternal inactivation throughout the developed female body (25).

Through random selection, most females will express the maternal or paternal X chromosome evenly. Females heterozygous for pathogenic variants will therefore express that variant in only half of their cells on average, compared to males who express the variant in all cells. In this way, XCI affords female resilience to X-linked diseases (26). By chance, however, a subset of females will demonstrate skewed expression to either allele (26), influencing disease severity by influencing the pathological burden within cells (26, 27). Pathogenic ALS/FTD-causing variants in *UBQLN2* likely cause gain of toxic function and loss of homeostatic functions (28). Thus, while females heterozygous for pathogenic *UBQLN2* variants are typically protected from these gain and loss of function effects compared to males by expressing the pathogenic variant in fewer cells, skew towards expression of the mutant allele reduces protection while skew towards the wildtype augments it.

Heterozygous females constitute the majority of reported *UBQLN2*-linked ALS/FTD cases (2) and yet are not represented by current disease models (6-13, 15, 19-21, 29-33). Previous models either recapitulate the toxic gain of function of pathogenic variants from the endogenous X-chromosome locus in hemizygous males (18, 33, 34) or overexpress the mutant gene from a transgenic locus (6-21). Modelling of *UBQLN2* loss of function is limited to complete knockdown of the gene (6, 8, 10, 12, 32, 35). Comparing models of heterozygous females that recapitulate the mosaic nature of *UBQLN2* expression from either the wildtype or mutant allele, to models with overall expression skewed towards either allele will inform our understanding of XCI in disease resilience in *UBQLN2*-linked ALS/FTD. Yet no such models have been described, and their establishment is limited by accurate measurement of the expression of each allele, which differ by a single base.

This paper describes the first characterisation of fibroblasts and iPSCs derived from females heterozygous for a pathogenic ALS/FTD-linked variant in *UBQLN2,* and an allele-selective qPCR (AS-qPCR) method for quantifying the proportion of mutant and wildtype allele that is expressed within a heterozygous population of cells. Together, this AS-qPCR assay and the assembly of a set of fibroblast and iPSC lines characterised for allelic expression will enable assessment of the influence of XCI and skew in female *UBQLN2* p.T487I-linked ALS/FTD and provide a model system for therapeutic development.

## Methods

### Participants and ethics

This study was approved by the Health and Disability Ethics Committee of New Zealand (HDEC approval 19/CEN/7). All participants provided informed written consent after telephone consultation with a genetic counsellor. All family and participant IDs used within this study are non-identifying and known only to select research members.

### Cell culture

#### Donor-derived fibroblast culture

Human dermal fibroblast (HDF) cultures were established at the University of Auckland, or by international collaborators local to the participant, by explant culture of skin punch biopsy tissue (36) (**Table S1**). Cultures were maintained in HDF growth medium (DMEM supplemented with 20% fetal bovine serum, 100 U/mL penicillin, 100 μg/mL streptomycin) and passaged using 0.25% Trypsin/EDTA in T75 flasks. Cells were maintained at 37 °C, 5% CO2 in a water-jacketed incubator to a maximum of 10 passages. All reagents were purchased from Life Technologies Ltd. unless stated otherwise.

#### iPSC generation and culture

Frozen vials of three HDF lines from females heterozygous for the p.T487I variant (Fibroblast lines 1, 2, 3) were delivered to StemCore (University of Queensland, Australia) on dry ice for reprogramming. Lines were reprogrammed using the Cytotune™-iPS 2.0 Sendai Reprogramming kit (Thermo Fisher Scientific) and twelve iPSC clones isolated per line (total 36 clones). Lines were maintained in mTeSR™ Plus medium (StemCell Technologies).

Following *UBQLN2* allelic skew quantification, the pluripotency and genetic stability of 12 selected clones were determined by StemCore by immunofluorescence for Oct4 (Abcam, AB283741, 1:200) and Nanog (Abcam, AB195018, 1:200), **Figure S1**), and the KaryoStat™ assay (Applied Biosystems, **Table S2, Table S3**). Lines were cryopreserved and vials delivered to the University of Auckland on dry ice.

### RNA extraction and cDNA synthesis

#### HDFs

HDF cells were plated at 80,000 cells/well in Nunclon™ 6-well plates and upon reaching 80% confluency were harvested for RNA extraction. Cells were washed with 1x ice-cold phosphate-buffered saline (PBS) and RNA extracted using the RNAqueous®-Micro Total RNA isolation Kit (Ambion) as per manufacturer’s instructions. RNA yield and purity were quantified using a Nanodrop spectrophotometer. RNA was stored at -80 °C prior to cDNA synthesis from 1 µg of RNA using the SuperScript IV Synthesis Kit as per manufacturer’s instructions.

#### iPSCs

iPSC RNA isolation and cDNA synthesis were performed by StemCore. RNA isolation was performed using the PureLink RNA Mini Kit (Ambion) and cDNA synthesis using the RevertAidFirst Strand cDNA Synthesis Kit (Thermo Fisher Scientific).

### *UBQLN2* allelic skew quantification

#### Sanger sequencing

Sanger sequencing was conducted by StemCore. An amplicon spanning the p.T487I variant site was generated from cDNA using primers as per **Table S6**. Amplicons were excised following gel electrophoresis and extracted using MinElute Gel extraction Kit (Qiagen). Samples were sent to the Australian Genome Research Facility (Brisbane, Australia) for Sanger sequencing (primers in **Table S6).** Chromatogram trace files (.ab1) and cDNA were sent to Auckland for quantification and Illumina and qPCR workflows, respectively. To semi-quantify *UBQLN2* allelic skew of iPSCs by Sanger sequencing, chromatogram files were read into R(v4.2.1) using the SangerseqR package (37). Semi quantification of the chromatogram at the p.T487I variant site was performed using the traceMatrix function.

#### Illumina amplicon sequencing

To quantify *UBQLN2* allelic skew of HDFs and iPSCs by Illumina sequencing, HDF cDNA generated in house and iPSC cDNA delivered by StemCore underwent tagged amplicon enrichment of *UBQLN2* by PCR using primers tagged with Illumina adapters (**Table S6**). Reactions used Phusion HF DNA polymerase (Thermo Fisher Scientific) following manufacturer’s instructions. *UBQLN2* amplicon enrichment was confirmed by gel electrophoresis and amplicons were purified using AmpureXP beads (Beckman Coulter) at a bead ratio of 0.9. Sequencing was performed by Auckland Genomics on an Illumina MiSeq using a 300 V2 Nano (2 x 150 bp) kit. Raw FASTQ files were provided and imported into R using the microseq package (38). Reads were trimmed to the p.T487I (c.1460C>T) variant site and calls with a Q score of ≥ 30 were excluded.

#### UBQLN2 allele-selective qPCR

An allele-selective qPCR (AS-qPCR) assay was developed to quantify relative expression of the *UBQLN2* WT and p.T487I alleles in HDFs and iPSCs. First, plasmid DNA (pDNA) was isolated to use as standards (pDEST-Myc *UBQLN2* WT and pDEST-Myc *UBQLN2* p.T487I) using the Nucleospin Plasmid kit (Macherey-Nagel) following manufacturer’s instructions. Transgene integrity was confirmed through Sanger sequencing.

Next, primers targeting the p.T487I variant site were designed in the reverse (sense-targeting) direction (**Figure S2**) positioning the p.T487I site at the 3’ terminal base (39-43). These primers were paired with allele-agnostic forward primers with similar predicted annealing temperatures (**Table S4**) but carrying no mismatches. Variant-targeting primers varied in length and either included or omitted a C:C mismatch located two bases upstream of their 3’ end to promote allelic selectivity (39, 40) while reducing inter-primer variability (39) and reducing off-target allele thermodynamic stability (44). Potential allele-selective primer pairs were screened at 65 °C using 1068 pg of pDEST-Myc *UBQLN2* WT and p.T487I pDNA (**Figure S3**).

All AS-qPCR reactions were conducted using a LightCycler® 480 II (Roche). The Platinum SYBR Green qPCR SuperMix-UDG with Rox was utilised per manufacturer’s instructions. qPCR reaction conditions are described in **Table S5**. Ct values were calculated with the LightCycler® 480 II software using 2nd Derivative Maximum Method. Amplification and melt curve traces were plotted using the ggplot2 package (45) in R. Of the primer pairs tested, Wildtype Reverse 4 and Mutant Reverse 4 with their respective forward primers produced robust and selective amplification of their respective targets across the entire range of allelic ratios and were therefore selected for the optimised assay.

To account for off-target amplification by the chosen and indeed all tested primer pairs due to the presence of a nearly identical template, a standard curve was generated using *UBQLN2* WT and p.T487I pDNA combined at varying ratios at a set total amount of template (46-50). First however, it was investigated whether a single standard curve could quantify *UBQLN2* in cDNA from both HDFs and iPSCs, by comparing their relative expression of *UBQLN2*.

RNA seq expression data was accessed from online databases. HDF transcript counts were accessed through GTEx analysis V8 dbGaP phs000424.v8.p2 (https://www.gtexportal.org/home/index.html) and “gene_tpm_cells_cultured_fibroblasts.gct” downloaded. *UBQLN2* and *TARDBP* counts were selected via ENSEMBL IDs ENSG00000188021.8 and ENSG00000120948.16 respectively. iPSC transcript counts quantified in transcripts per million (TPM) were accessed through Stemformatics (https://www.stemformatics.org/) and iPSCs derived from fibroblasts selected and individual datasets downloaded. Datasets were collated, *UBQLN2* and *TARDBP* counts selected using the above ENSEMBL IDs, and disease state set to normal. Data was managed and visualized using R and visualized with the ggplot2 package. Expression levels in both lines were low but comparable (**Figure S4A**) such that a single standard curve could be used for AS-qPCR of both cell types.

Next, the amount of pDNA per reaction required to generate a standard curve was determined. pDEST-Myc *UBQLN2* WT pDNA was serially diluted until an allele-agnostic *UBQLN2* primer pair (**Table S6**) gave a Ct value equal to that using 10 ng of male control HDF cDNA (**Figure S4 B**). The amount of template for standard curve generation was determined to be 1.04 pg of pDNA/reaction. Therefore, a final standard curve of Ct values was generated using the hit primer sets across the range of allelic ratios using 1.04 pg total template / reaction (38-42) (**Figure S3 C**). Finally, AS-qPCR was benchmarked against the allele-agnostic *UBQLN2* primer pair.

### Quantification of escape from X chromosome inactivation

#### Immunocytochemistry

iPSC lines were plated for immunocytochemistry for ubiquilin 2 and Barr body marker H3K27me3. Cells were plated at 5,000 cells/well into a Matrigel® (Corning) coated Nunclon™ 96-well plate in mTesR™ Plus supplemented with RevitaCell™ (Gibco) to aid cell survival. Media was replaced without RevitaCell™ every 24 h until ∼50% cell confluency was reached. An additional male wildtype iPSC line (KOLF2.1J, RRID:CVCL_B5P3, The Jackson Laboratory) was cultured in StemFlex™ media (Gibco) and underwent immunocytochemistry as a negative control for the female iPSC clones. Passage number was kept below 7 to minimise genetic drift across passages.

Cells were fixed in 4% paraformaldehyde for 15 min and washed in 1x PBS containing 0.2% (v/v) Triton X-100 (PBS-T). Cells were incubated with mouse monoclonal ubiquilin 2 antibody (Novus Biologicals, NBP2-25164) and rabbit monoclonal H3K27me3 antibody (Abcam, ab192985) each diluted in 1:500 in PBS-T with 1% normal goat serum (goat immunobuffer) overnight at 4 °C. Cells were then washed 2x with PBS-T, before incubation for 5 h at room temperature with goat anti-mouse AlexaFluor^TM^ Plus 488 and goat anti-rabbit AlexaFluor^TM^ 594 both diluted 1:500 in goat immunobuffer with 10 µg/mL Hoechst 33358.

Images of cells were taken on a PerkinElmer Operetta high-content analysis system at 20x NA magnification using settings DAPI (Ex/Em 355 – 385/410 – 480 nm), AlexaFluor 488 (Ex/Em 460 – 490/550 – 550 nm) and Alexafluor 594 (Ex/Em 530 – 560/590 – 640 nm). Images were captured as .tif files and uploaded to the CellProfiler v4.7.2 image analysis tool (51). In brief, cell nuclei and H3K27me3 nuclear puncta were enhanced through the modules HistogramEqualization (52) and EnhanceOrSuppressObjects (51) and used to delineate individual Barr bodies using IdentifyPrimaryObjects. Local minima closer than 4 pixels were suppressed. The perinuclear mean ubiquilin 2 labelling intensity was identified per cell using the ExpandOrShrinkObjects (5 pixels), MaskImage and MeasureObjectIntensity modules. The number of H3K27me3 puncta per cell was calculated using the RelateObjects module. Per-cell measurements were imported into R(v4.5.2). Ubiquilin 2 signal intensity was normalised to the no primary antibody control of each line. H3K27me3 puncta (Barr bodies) per cell were classified as “0”, “1”, or “>1”.

#### Statistical analyses

To analyse escape from X inactivation, 6,000 iPS cells were randomly subsampled per line and a linear mixed effects regression model fitted using the lme4 package (53). Fixed effects were Barr body number with random intercepts specified for the imaging hierarchy of fields of view, wells, and cell lines. Type III analysis of variance (ANOVA) of fixed effects was conducted using Satterthwaite’s approximation for degrees of freedom through the emmeans package (54), and pairwise comparisons performed. Standardised effect sizes (Cohen’s *d*) between fixed effects were calculated using the eff_size function with residual standard deviation and degrees of freedom inherited from the model. Effect size biological significance thresholds were small (< 0.2), medium (< 0.6), and large (≥ 0.8), as referenced in **Figure 2**.

To analyse Barr body number and non-dominant allele expression correlation, the allelic ratios of 12 chosen iPSC lines, as measured by AS-qPCR, Sanger sequencing, or Illumina amplicon-based sequencing, were correlated with the observed proportion of cells exhibiting a single Barr body. A linear mixed effects model was fitted for each allelic measurement technique using restricted maximum likelihood with the slope of the regression being the fixed effect, and random intercepts for each parental line of each clone to account for parent-dependent variability. Summary statistics were calculated using the summary function, and marginal coefficient of determination calculated using the performance function.

All statistical tests were two-sided, and significance was set at α = 0.05.

## Results

### Allelic skew was variable in heterozygous female fibroblast lines but strong in resultant clonal iPSCs

Dermal fibroblast lines established from four females heterozygous for a *UBQLN2* c.1460C>T (p.T487I) variant expressed both *UBQLN2* alleles, although the relative contribution of each allele varied between individuals, consistent with mosaic populations of cells from XCI (**Figure 1 A**). The Fibroblast 1 line showed near-equal expression of mutant and wildtype alleles while the remaining heterozygous lines demonstrated varying degrees of skew toward the wildtype allele. Homozygous and hemizygous wildtype controls (two donors each) expressed only the wildtype allele.

**Figure 1.**
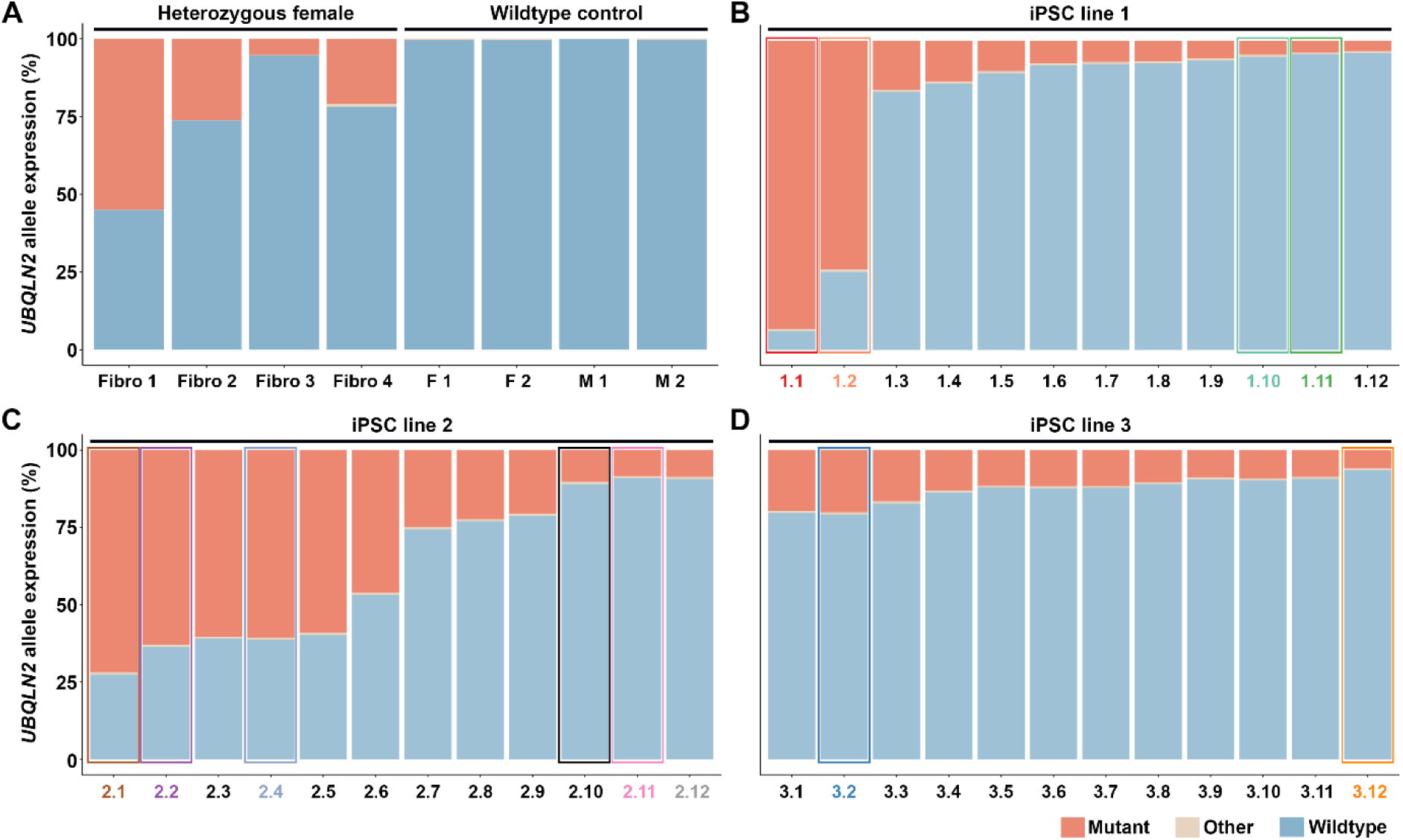
Allelic skew in fibroblast lines heterozygous for *UBQLN2* p.T487I, and their derived iPSC clones. **(A)** Relative expression of mutant and wildtype *UBQLN2* alleles in four fibroblast lines from females heterozygous for *UBQLN2* p.T487I alongside wildtype controls. **(B-D).** Allelic skew in 12 clonal iPSCs lines derived from lines Fibroblast 1 (**B**), Fibroblast 2 (**C**), and Fibroblast 3 (**D**). Allelic expression was quantified by Illumina-based amplicon sequencing. Clones selected for subsequent analyses boxed and colour-coded. Abbreviations: F, female; M, male.

iPSC reprogramming and clonal isolation of three of the fibroblast lines (1, 2, and 3) produced a total of 36 clones (12 per line) that were sequenced for allelic expression (**Figure 1 B-D**). The fourth fibroblast line was not reprogrammed. Many clones from the Fibroblast 1 line demonstrated comparatively biallelic expression while the Fibroblast 2 line skewed strongly toward either the mutant, or wildtype allele. Clones derived from the Fibroblast 3 line skewed strongly toward the wildtype allele.

Of the 36 clones sequenced, 12 clones were selected (2-6 clones per parent fibroblast line) to model a range of allelic skews. These included comparatively biallelically expressing clones and strongly skewed clones to further investigate the association between XCI persistence and ubiquilin 2 expression.

### Clonal iPSC lines demonstrated variable Barr body number and Barr body absence increased total ubiquilin 2 expression

The persistence of XCI is critical to the utility of the selected iPSC clones as models of heterozygous of *UBQLN2* linked p.T487I ALS/FTD. To determine XCI integrity, Barr body number was visualised and quantified in the 12 selected iPSC clones and compared with a male iPSC line used as a negative control (**Figure 2 A and B**). Most cells within each variant-carrying clone produced a single H3K27me3 punctum, representing a Barr body, consistent with maintenance of XCI, however all clones also contained a subset of cells devoid of detectable Barr bodies (**Figure 2 C**). Unexpectedly, subsets of cells within some clones were also identified that carried more than one Barr body, such as in Clone 2.4 (**Figure 2 A and C**). As expected, no Barr bodies were identified in the male control line (**Figure 2 A and C**).

**Figure 2.**
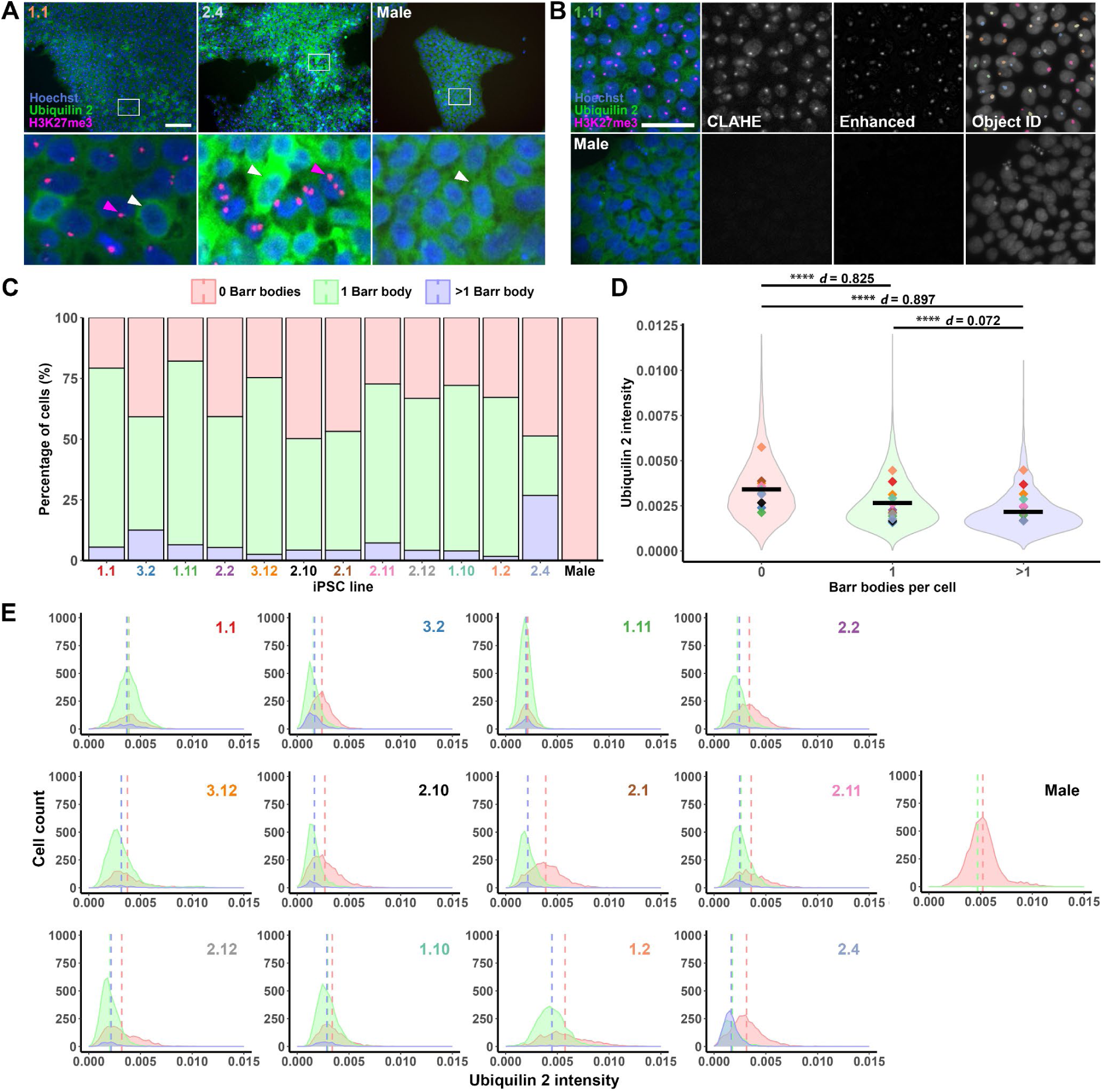
Clonal iPSC lines exhibit variable Barr body number and loss of Barr bodies is associated with increased ubiquilin 2 expression. **(A)** Representative images of a clone predominantly expressing a single Barr body (Clone 1.1), a clone predominantly expressing either 0 or >1 Barr body (Clone 2.4), and the male control line expressing 0 Barr bodies. White arrows indicate representative cells with no Barr body. Purple arrows denote H3K27me3-positive nuclear puncta corresponding to Barr bodies. Scale bar is 100 microns. **(B)** Representative images of H3K27me3 puncta enhancement using Contrast Limited Adaptive Histogram Equalization (CLAHE) and subsequent object identification in a female line (Clone 1.11) and male negative control. Scale bar is 50 microns. **(C)** Percentage of cells expressing 0, 1, or >1 Barr bodies across the 12 selected female iPSC clones compared to a male control line. **(D)** Violin plot of pooled ubiquilin 2 intensities from the 12 female iPSC clones grouped by Barr bodies per cell. Statistical significance was determined using a linear mixed-effects model, **** = p < 0.0001, d = effect size. Coloured diamonds represent clone mean values; black bars represent group mean. **(E)** Histograms of ubiquilin 2 staining intensity of each line by Barr body number. Red, green and blue vertical lines represent mean ubiquilin 2 staining intensity for populations of cells with 0, 1, or >1 Barr body, respectively for each line. Data shown correspond to the 0.5 - 99.5^th^ percentile of 6000 randomly subsampled cells per line to ensure balanced sampling and minimise the influence of outliers.

To determine the effect of Barr body number on ubiquilin 2 expression, ubiquilin 2 staining intensity was calculated per cell and stratified by Barr body number, either in all 12 clones pooled (**Figure 2 D)** or in individual clones (**Figure 2 E**). Cells with no Barr body showed a significant increase in the intensity of total ubiquilin 2 compared to cells carrying 1 or more Barr bodies, with a large effect size (**Figure 2 D,** p < 0.0001). The presence of more than 1 Barr body was associated with significantly less intensity of ubiquilin 2 compared to cells with 1 Barr body, but with a small effect size.

To further characterise the clones, all 12 underwent karyotyping using the CytoScan® array to assess the presence of copy number variants (CNVs) (i.e. InDels) and loss of heterozygosity (LOH) across the genome. Karyotyping of the lines did not identify any CNVs on the X chromosome in any line aside from clone 3.2 (**Table S2**). A consistent LOH genotype was found in all lines near the centromere between cytobands ∼q11-12 and q13 - 21 (**Table S3**), distinct to the locus of *UBQLN2* (p11.21) (55).

Taken together, the 12 selected iPSC clones heterozygous for the ALS/FTD-linked p.T487I variant demonstrated variable retention of a single Barr body between clones with Barr body absence increasing total ubiquilin 2 protein. This finding informs which clone/s best retain the disease-relevant phenomenon of X chromosome inactivation.

### Validation of allelic skew in heterozygous female clonal iPSCs using a rapid and affordable allele-selective qPCR method

While Illumina-based amplicon sequencing enables accurate quantification of single nucleotide variants, the process is both time-consuming and costly. Allele-selective qPCR (AS-qPCR) offers a high-throughput solution to quantify the skew toward *UBQLN2* wildtype or p.T487I alleles in a bulk population of cells.

Primers with confirmed selectivity for either the p.T487I mutant, the wildtype allele, or allele-agnostic primers (**Figure S3**), were tested against wildtype and mutant plasmid DNA (pDNA), combined across a range of ratios at an optimised amount of total template (**Figure S4 B**), to generate a standard curve representing the full spectrum of potential *UBQLN2* skew (**Figure S4 C**) (46, 47, 50). Efficient amplification was observed across all allelic ratios using the wildtype primer, including in the absence of any wildtype template, indicating this primer is wildtype *selective* not specific. However, while efficient amplification with the mutant-selective primer was observed with as little as 3.125% mutant allele, when only the wildtype allele was present using 1.04 pg of pDNA, inefficient amplification was observed and a default Ct value of 40 was assigned to the standard curve, indicating this primer is fully mutant specific. ΔCt values were calculated using Equation 1 and a piecewise cubic spline regression with knots at 6.25 and 93.5% was fitted to generate the standard curve (**Figure 3 A**) (56, 57).

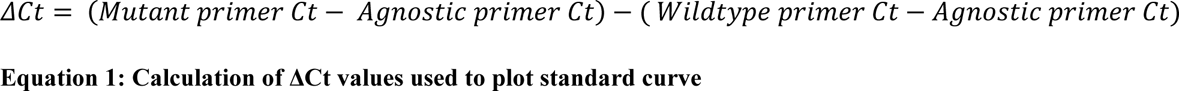

**Figure 3.**
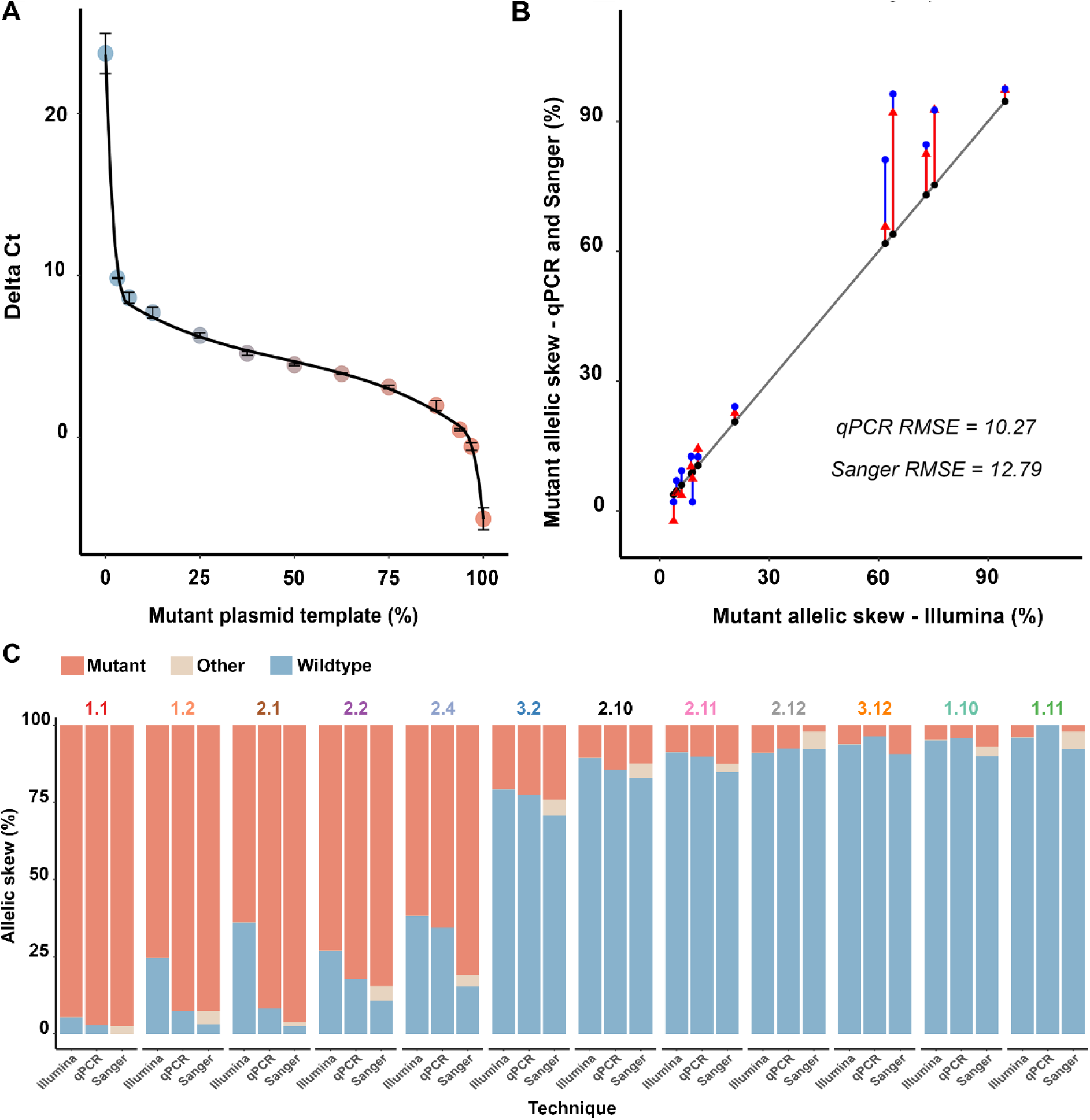
Validation of AS-qPCR assay against Sanger semi-quantification and Illumina amplicon sequencing in iPSC clones heterozygous for *UBQLN2* p.T487I. **(A)** Standard curve relating AS-qPCR Delta CT values to known mutant:wildtype template ratios. Data represent mean +/- SEM from two independent biological replicates and were fitted using a piecewise cubic spline regression with knots at 6.25% and 93.5% mutant plasmid template. **(B)** Mutant allelic load determined by Illumina amplicon sequencing (black dots) compared to estimates obtained by AS-qPCR (red triangles) and Sanger chromatogram quantification (blue circles). Root-mean square error (RMSE) for AS-qPCR and Sanger are shown relative to Illumina sequencing. (**C**) Comparison of allelic skew across clonal iPSC lines as measured by AS-qPCR, Illumina amplicon, and Sanger chromatogram quantification.

Using this standard curve, the accuracy of the AS-qPCR assay was benchmarked using cDNA from the 12 iPSC clones heterozygous for the p.T487I variant against Illumina sequencing and semi-quantification of Sanger chromatograms (**Figure 3 B and C**). The AS-qPCR assay accuracy was comparable to Sanger sequencing semi-quantification (RSME= 12.79) and Illumina sequencing (RSME= 10.27) across all samples.

Taken together, the allelic skew towards wildtype or p.T487I transcripts could be measured with high accuracy by the AS-qPCR assay developed here, confirming that while most clonal iPSC populations demonstrated strong allelic skew, a subsection produced biallelic expression.

### Abnormal Barr body number correlates with biallelic expression from iPSC lines

AS-qPCR, Sanger chromatogram quantification, and Illumina sequencing all confirmed that most clones were strongly skewed toward expression of a single allele, as expected given XCI is inherited by daughter cells in a clonal population. However, a subset of clones exhibited varying degrees of non-dominant allele expression, suggestive of partial loss of allelic skew. This may reflect XCI erosion (58-61), or abnormal Barr body formation as observed in Clone 2.4 (**Figure 2**).

To investigate this possibility, non-dominant allele expression predicted by each technique was correlated with the proportion of cells of each iPSC clone carrying the expected single Barr body (**Figure 4**). The proportion of cells carrying a single Barr body was significantly inversely correlated with expression of the non-dominant allele in clonal populations when measured by qPCR (R^2^= 0.90, p< 0.0001), Sanger sequencing (R^2^= 0.37, p= 0.0216), or Illumina amplicon sequencing (R^2^= 0.54, p= 0.0049), albeit with varying goodness of fit of the linear model.

**Figure 4.**
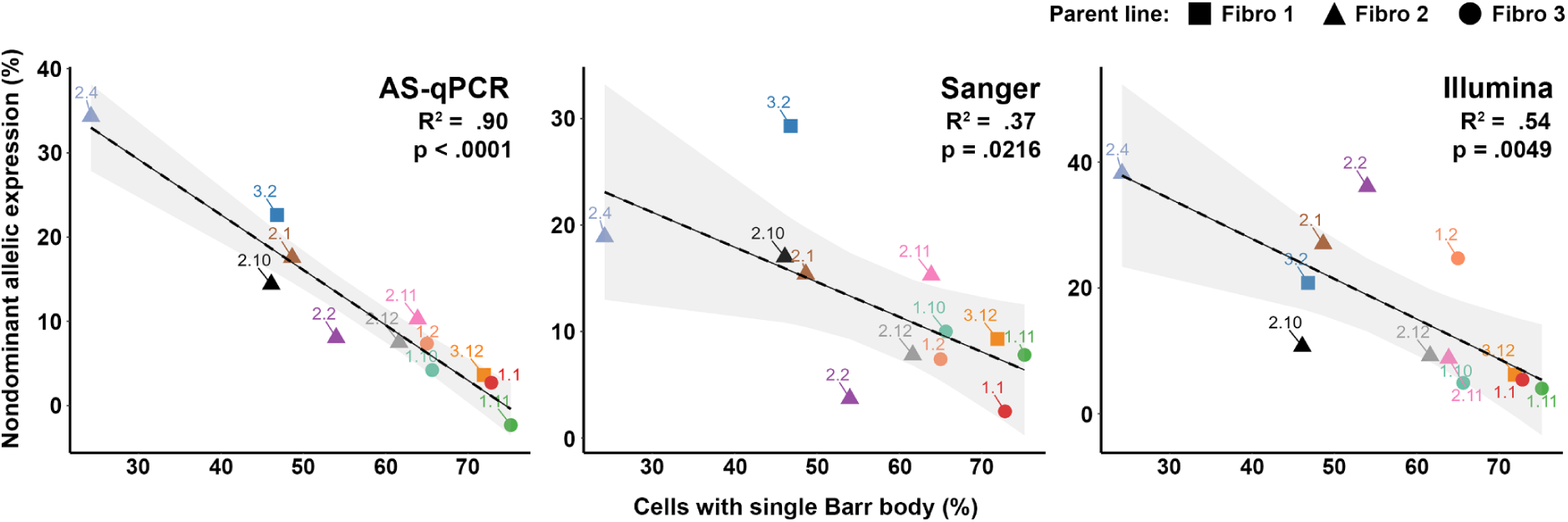
Correlation between allelic skew and Barr body number across iPSC clones using three independent quantification methods. Non-dominant allelic expression measured by AS-qPCR, Illumina amplicon sequencing, and Sanger chromatogram quantification was correlated with the percentage of cells containing a single Barr body within each iPSC clone. Shaded regions indicate the 95% confidence intervals of the fitted regression lines. Insets display marginal coefficient of determination (R^2^) and corresponding p values. Different point shapes denote parental fibroblast line from which each clone was derived.

This indicates that AS-qPCR had greater sensitivity to identify the correlation between allelic expression and escape from XCI than Illumina sequencing or Sanger chromatogram semi-quantification, effectively finding that biallelic expression/ loss of skew in bulk populations of iPSCs which should be clonal is due to abnormal XCI dynamics.

## Discussion

Consistent with other X chromosome-linked disorders (26), *UBQLN2*-linked ALS/FTD demonstrates sex-dependent heterogeneity in age at onset and death (2, 4), which is likely due to stochastic skewing of XCI between tissues. Despite heterozygous females constituting most published cases (2), current models of *UBQLN2-*linked ALS/FTD have not examined the effect of variable XCI-mediated pathogenic variant skew on the pathophenotype (6-13, 15, 19-21, 29-33). To do that, one would ideally compare pathophenotypes between mixtures of isogenic iPSC lines expressing mutant or wildtype *UBQLN2* from the endogenous X-chromosome locus, with cells combined at various ratios to model the mosaicism of female tissues. Such a model would also enable high-throughput testing of therapies to selectively deplete the mutant allele while retaining wildtype ubiquilin 2 expression. However, female *UBQLN2* iPSCs have not been reported and iPSCs are known to suffer erosion of XCI that would confound analysis of its role. Moreover, gold-standard methods of single nucleotide variant quantification to validate the model and the effect of the therapy remain laborious and expensive. Together, these limitations reduce the reliability of iPSC-derived lines as a reporter of the role of XCI in female resilience and hinder the validation of allele-selective therapies for *UBQLN2*-linked ALS/FTD.

Here we characterised 12 iPSC clones derived from 3 heterozygous carriers of the ALS/FTD-linked p.T487I variant in *UBQLN2* and demonstrated the development of an accurate, inexpensive, and efficient allele-selective qPCR assay to measure allelic skew in bulk samples.

### Erosion of XCI as a factor in the selection of iPSC clones for modelling *UBQLN2*-linked ALS/FTD and other X-linked disorders

The allelic skew of four fibroblast lines and 36 resultant iPSC clones was quantified through amplicon-based Illumina sequencing. As predicted, fibroblast lines demonstrated variable skew toward either allele. Deviations from the ‘normal’ 50:50 allelic expression have been reported across tissues (62, 63) in 10 - 50% of females (64-68), with fibroblasts also demonstrating heterogeneity in the fidelity of XCI (69). In contrast, most iPSC clones demonstrated strong skew as expected for lines derived from a single cell with heritable XCI (60, 70-72). However, a subset of cells in certain iPSC clones demonstrated biallelic expression, indicating a degree of escape from or erosion of XCI.

Erosion of XCI was assessed on a per-cell basis by examining H3K27me3 as a Barr body marker in twelve selected iPSC clones in which *UBQLN2* expression was also quantified. We found that while most cells from most iPSC clones produced a single Barr body, a small proportion of cells from certain lines harboured more than 1 Barr body, exemplified by iPSC Clone 2.4. This could be explained by karyotypic changes (e.g. trisomy of the X chromosome (73-75), although this was not borne out by copy number analysis), or by image analysis artefact for most clones. Importantly, there was also a subpopulation of cells without a Barr body which expressed approximately twice as much ubiquilin 2. XCI erosion is a well-documented phenomenon in stem cell lines (58-61, 76). It is hypothesised to be dependent upon the presence of lithium chloride in cell media (including mTESR Plus, in which these lines were cultured) (61). Our findings indicate that, consistent with previous literature (58-60), stripping of H3K27me3 is a marker of the erosion of X inactivation. Furthermore, we have shown that *UBQLN2* became functionally biallelically expressed in certain iPSC clones to variable degrees which is likely to be due to erosion of XCI (77, 78).

Together, these analyses indicated which patient-derived iPSC clones retained XCI and expressed *UBQLN2* from a single allele, as occurs physiologically in humans (78), and are thus the most useful for disease modelling.

### Development of an allele-selective qPCR assay and the relationship between allelic skew and Barr body number

Although we had determined *UBQLN2* allelic expression of the 36 iPSC clones (and their parent fibroblast lines) using Illumina amplicon-based sequencing, we sought a rapid, inexpensive alternative in the form of a novel allele-selective qPCR (AS-qPCR) assay.

Amplicon-based NGS technologies (such as Illumina sequencing) are often seen as a gold-standard to quantify the expression of variant alleles (79, 80). However, amplicon generation prior to base calling can introduce error (81-83). The HUMARA assay, which examines the polymorphic expansion of the human androgen receptor gene (84), has been considered a gold standard to measure overall XCI skew, but is uninformative in the 10-25% of females homozygous for the expansion (66, 85, 86), nor directly informs allelic skew at the locus of interest (pertinent when measuring gene-specific escape from XCI). Long-read sequencing of the transcriptome (87, 88) is the most comparable to AS-qPCR for measuring allelic skew but is costly and requires high quality RNA input (89). CRISPR-enriched Long-read sequencing and methylation analysis of DNA negates the requirement for expression in the sampled tissue (90, 91) but cost and labour intensive sample preparation issues remain (89). We sought an inexpensive assay that would best enable characterisation of *UBQLN2* skew in iPSC lines at baseline, as described here, but also across differentiation and in response to potential gene therapies in future work.

A novel allele-selective *UBQLN2* AS-qPCR assay was developed to measure allelic skew and benchmarked against Illumina sequencing and Sanger chromatogram semi-quantification using the same cDNA sample. qPCR quantification is frequently normalised to an internal control (e.g. ‘housekeeper’ gene) with the assumption that the target and reference primers remain equally efficient across a range of concentrations (92). For AS-qPCR, the presence of varying amounts of a near identical template makes this assumption difficult to uphold. Therefore, a standard curve of allelic skews was employed, as recommended previously (47, 50, 93). AS-qPCR primer design also drew on previously described principles, with maximal allele selectivity conferred by positioning the base pair mismatch at the primer’s 3’ terminus (39-43). This assay confirmed that the increased ubiquilin 2 expression associated with erosion of XCI marker H3K27me in *UBQLN2* iPSC clones such as Clone 2.4 was due to abnormal Barr body dynamics. In contrast, iPSC clones such as 1.1 (mutant skew) and 1.11 (wildtype skew) which had the greatest proportion of cells with a single Barr body showed the strongest skew towards the dominant allele. These latter clones show XCI persistence, as occurs in adult female tissues, and therefore represent ideal models to further examine the effect of XCI in *UBQLN2*-linked ALS/FTD.

Overall, correlating Barr body number with AS-qPCR- and orthogonal assay- validated non-dominant allele load (i.e. biallelic expression) enabled the identification of heterozygous carrier-derived iPSC clones which have largely preserved XCI and validated an assay for high-throughput quantification of the allelic load from bulk-cell populations. Together, these findings provide a foundation for future work to explore the under-researched role of XCI in female *UBQLN2*-linked ALS/FTD.

## Conclusion

The effect of X chromosome inactivation (XCI) in *UBQLN2*-linked ALS/FTD is not represented in existing disease models, despite heterozygous females comprising the majority of affected carriers. In this study, twelve iPSC clones were selected from 3 females heterozygous for the pathogenic *UBQLN2* p.T487I variant. Immunocytochemistry confirmed the preservation of XCI in most clones although a subset exhibited evidence of XCI erosion accompanied by increased in ubiquilin 2 protein expression. A novel allele-selective qPCR assay further revealed a strong correlation between the proportion of cells with XCI erosion and increased expression of the non-dominant *UBQLN2* allele. These analyses identified iPSC clones that most faithfully preserve the disease-relevant phenomenon of XCI. Together with the newly developed AS-qPCR assay, these models provide a foundation for future studies investigating the role of XCI and the development of therapeutic strategies in female *UBQLN2*-linked ALS/FTD.

