## Supplementary material for "Quantification of X-chromosome inactivation in fibroblast and iPSC models of *UBQLN2* ALS/FTD using allele-selective qPCR"

Table S1 Biological case details for human dermal fibroblast and their resulting iPSC lines

| **Line ID** | **Cell types** | **Genotype** | **Sex** |
| --- | --- | --- | --- |
| Fibro 1 | Fibroblast and iPSC | Heterozygous c.1460C>T, p.T4871 | Female |
| Fibro 2 | Fibroblast and iPSC | Heterozygous c.1460C>T, p.T4871 | Female |
| Fibro 3 | Fibroblast and iPSC | Heterozygous c.1460C>T, p.T4871 | Female |
| Fibro 4 | Fibroblast only | Heterozygous c.1460C>T, p.T487I | Female |
| M 1 | Fibroblast only | Hemizygous WT control | Male |
| M 2 | Fibroblast only | Hemizygous WT control | Male |
| F 1 | Fibroblast only | Homozygous WT control | Female |
| F 2 | Fibroblast only | Homozygous WT control | Female |

Note. Age at donation and donor phenotype are omitted to avoid participant (self-)identification

Table S2 Heterozygous iPSC CNV InDel report by CytoScan® Optima Assay

| **Identification** | | | **CNV InDels** | | | | | |
| --- | --- | --- | --- | --- | --- | --- | --- | --- |
| **Clone** | **Parent** | **Sex** | **Aberration type** | **Copy number** | **Chromosome** | **Cytoband** | **Marker boundaries** | **Size** |
| 1.10 | Fibro 1 | Female | Loss | 1 | chr11 | q14.1 | 80,855,353-81,318,735 | 463 |
| 1.11 | Fibro 1 | Female | Loss | 1 | chr11 | q14.1 | 80,855,353-81,318,735 | 463 |
| 2.2 | Fibro 2 | Female | Gain | 3 | ch1 | q24.3-q25.1 | 172,330,984-174,339,289 | 2,008 |
| 3.2 | Fibro 3 | Female | Gain | 3 | chrX | p22.32-p22.31 | 5,990,149-8,723,023 | 2,733 |
|  |  |  | Gain | 3 | chrX | p22.2-p22.13 | 15,783,833-17,720,899 | 1,937 |
|  |  |  | Gain | 3 | chrX | p22.12-p22.11 | 20,475,186-22,010,406 | 1,535 |
|  |  |  | Gain | 3 | chrX | q21.31-q22.1 | 91,734,047-99,574,559 | 7,841 |
|  |  |  | Gain | 3 | chrX | q22.2 | 103,466,033-104,224,546 | 759 |
|  |  |  | Gain | 3 | chrX | q23 | 111,294,514-116,187,254 | 4,893 |

Marker boundaries annotated to hg38. Size in kbp.

Table S3 Heterozygous iPSC CNV Loss of heterozygosity report by CytoScan® Optima Assay

| **Identification** | | | **Loss of heterozygosity** | | | |
| --- | --- | --- | --- | --- | --- | --- |
| **Clone** | **Parent line** | **Sex** | **Chromosome** | **Cytobands** | **Location** | **Size** |
| Control | NA | Male | chrX | p22.33-p11.1 | 2,686,633-58,311,457 | 55,625 |
|  |  |  | chrX | q11.1-q28 | 62,713,034-155,993,566 | 93,281 |
| 1.1 | Fibro 1 | Female | chX | q11.1-q12 | 62,713,034-67,862,682 | 5,150 |
|  |  |  | chrX | q13.1-q21.1 | 71,541,431-79,799,444 | 8,258 |
| 1.2 | Fibro 1 | Female | chrX | q11.1-q12 | 62,713,034-68,162,493 | 5,449 |
| 1.10 | Fibro 1 | Female | chrX | q11.1-q12 | 62,713,034-68,084,866 | 5,372 |
|  |  |  | chrX | q13.1-q21.1 | 71,541,431-79,259,214 | 7,718 |
| 1.11 | Fibro 1 | Female | chrX | q11.1-q12 | 62,713,034-68,084,866 | 5,372 |
|  |  |  | chrX | q13.1-q21.1 | 71,541,431-79,259,214 | 7,718 |
| 2.1 | Fibro 2 | Female | chrX | q11.1-q12 | 62,713,034- 68,151,635 | 5,439 |
|  |  |  | chrX | q13.1-q21.1 | 72,103,250-77,364,171 | 5,261 |
| 2.2 | Fibro 2 | Female | chrX | q11.1-q12 | 62,713,034-68,084,866 | 5,372 |
|  |  |  | chrX | q13.1-q21.1 | 71,386,358-77,364,171 | 5,978 |
| 2.4 | Fibro 2 | Female | chrX | q11.1-q12 | 62,713,034- 68,162,493 | 5,449 |
| 2.10 | Fibro 2 | Female | chrX | q11.1-q12 | 62,713,034- 68,084,866 | 5,372 |
|  |  |  | chrX | q13.1-q21.1 | 71,386,358-77,364,171 | 5,978 |
| 2.11 | Fibro 2 | Female | chr20 | q11.21-q11.23 | 32,404,488-37,585,513 | 5,181 |
|  |  |  | chrX | q11.1-q12 | 62,713,034-68,151,635 | 5,439 |
|  |  |  | chrX | q13.1-q21.1 | 71,386,358-77,452,259 | 6,066 |
| 2.12 | Fibro 2 | Female | chrX | q11.1-q12 | 62,713,034-68,151,635 | 5,439 |
|  |  |  | chrX | q13.1-q21.1 | 71,386,358-77,364,171 | 5,978 |
| 3.2 | Fibro 3 | Female | chrX | q11.1-q12 | 62,713,034-68,151,635 | 5,439 |
|  |  |  | chrX | q13.1-q21.1 | 71,386,358-77,364,171 | 5,978 |
| 3.12 | Fibro 3 | Female | chrX | q11.1-q12 | 62,713,034-68,162,493 | 5,449 |

Marker boundaries annotated to hg38. Size in kbp.

Table S4 Predicted Tm of allele-selective reverse primers and paired allele-agnostic forward primer against their target template

| **Forward primer** | **Predicted Tm ºC** | **Reverse Primer** | **Predicted Tm ºC** |
| --- | --- | --- | --- |
| Forward 1 | 56.0 | Wildtype Reverse 1 | 59.5 |
| Forward 5 | 62.7 | Wildtype Reverse 2 | 62.8 |
| Forward 5 | 62.7 | Wildtype Reverse 3 | 62.5 |
| Forward 6 | 67.3 | Wildtype Reverse 4 | 67.5 |
| Forward 6 | 67.3 | Wildtype Reverse 5 | 67.3 |
| Forward 6 | 67.3 | Wildtype Reverse 6 | 71.7 |
| Forward 2 | 57.7 | Mutant Reverse 1 | 58.5 |
| Forward 5 | 62.7 | Mutant Reverse 2 | 62.2 |
| Forward 4 | 61.2 | Mutant Reverse 3 | 51.9 |
| Forward 6 | 67.3 | Mutant Reverse 4 | 67.6 |
| Forward 6 | 67.3 | Mutant Reverse 5 | 58.7 |
| Forward 6 | 67.3 | Mutant Reverse 6 | 64.9 |

Primer sequences and genomic locations are outlined in Table S6 and Fig S2.

Table S5 qPCR reaction conditions for AS-qPCR

| **Temperature - time** | **Ramp speed** | **Cycle number** | **Step** |
| --- | --- | --- | --- |
| 95 °C – 10 sec | 4.8 °C/s | 1 | Initial denaturation |
| 95 °C – 15 sec | 4.8 °C/s |  | Denaturation |
| Variable – 1 min | 2.5 °C/s | 40 | Annealing |
| 72 °C – 15 sec | 4.8 °C/s |  | Extension |
| 95 °C – 15 sec | 4.8 °C/s | 1 | Melt curve |
| 63 °C – 1 min | 2.5 °C/s | 1 |  |
| 95 °C – 15 sec | 0.11 °C/s | 1 |  |

Table S6 Primer list

| **Purpose** | **Direction** | **Sequence (5' to 3')** |
| --- | --- | --- |
| Sanger amplicon generation | Forward | CAGCAGGGGCTACAGACATT |
| Sanger amplicon generation and sequencing | Reverse | CTATGGGGGTAAAAGGGACT |
| Illumina enrichment with adaptor tags | Forward | TCGTCGGCAGCGTCAGATGTGTATAAGAGACAGCAGCAGGGGCTACAGACATT |
| Illumina enrichment with adaptor tags | Reverse | GTCTCGTGGGCTCGGAGATGTGTATAAGAGACAGCTATGGGGGTAAAAGGGACT |
| Allele-selective qPCR (Forward 1) | Forward | AGACATTAGCCACTGAAGCA |
| Allele-selective qPCR (Forward 2) | Forward | TCAGCCATGTCAAACCCAA |
| Allele-selective qPCR (Forward 4) | Forward | AGCCATGTCAAACCCAAGAGC |
| Allele-selective qPCR (Forward 5) | Forward | TGAAGCACCTGGCCTGATTCC |
| Allele-selective qPCR (Forward 6) | Forward | GCCACTGAAGCACCTGGCCTGA |
| Allele-selective qPCR (Wildtype Reverse 1) | Reverse | TACAGGGCCTATAGCGG |
| Allele-selective qPCR (Wildtype Reverse 2) | Reverse | CCTACAGGGCCTATAGCGG |
| Allele-selective qPCR (Wildtype Reverse 3) | Reverse | CCTACAGGGCCTATACCGG |
| Allele-selective qPCR (Wildtype Reverse 4) | Reverse | GGCCTACAGGGCCTATAGCGG |
| Allele-selective qPCR (Wildtype Reverse 5) | Reverse | GGCCTACAGGGCCTATACCGG |
| Allele-selective qPCR (Wildtype Reverse 6) | Reverse | ACTGGGCCTACAGGGCCTATACCGG |
| Allele-selective qPCR (Mutant Reverse 1) | Reverse | TACAGGGCCTATAGCGA |
| Allele-selective qPCR (Mutant Reverse 2) | Reverse | CCTACAGGGCCTATAGCGA |
| Allele-selective qPCR (Mutant Reverse 3) | Reverse | CCTACAGGGCCTATACCGA |
| Allele-selective qPCR (Mutant Reverse 4) | Reverse | GGCCTACAGGGCCTATAGCGA |
| Allele-selective qPCR (Mutant Reverse 5) | Reverse | GGCCTACAGGGCCTATACCGA |
| Allele-selective qPCR (Mutant Reverse 6) | Reverse | ACTGGGCCTACAGGGCCTATACCGA |
| Allele-agnostic *UBQLN2* qPCR primer | Forward | AGACATTAGCCACTGAAGCA |
| Allele-agnostic *UBQLN2* qPCR primer | Reverse | TCAGCCATGTCAAACCCAA |

Underlined bases indicate Illumina tags

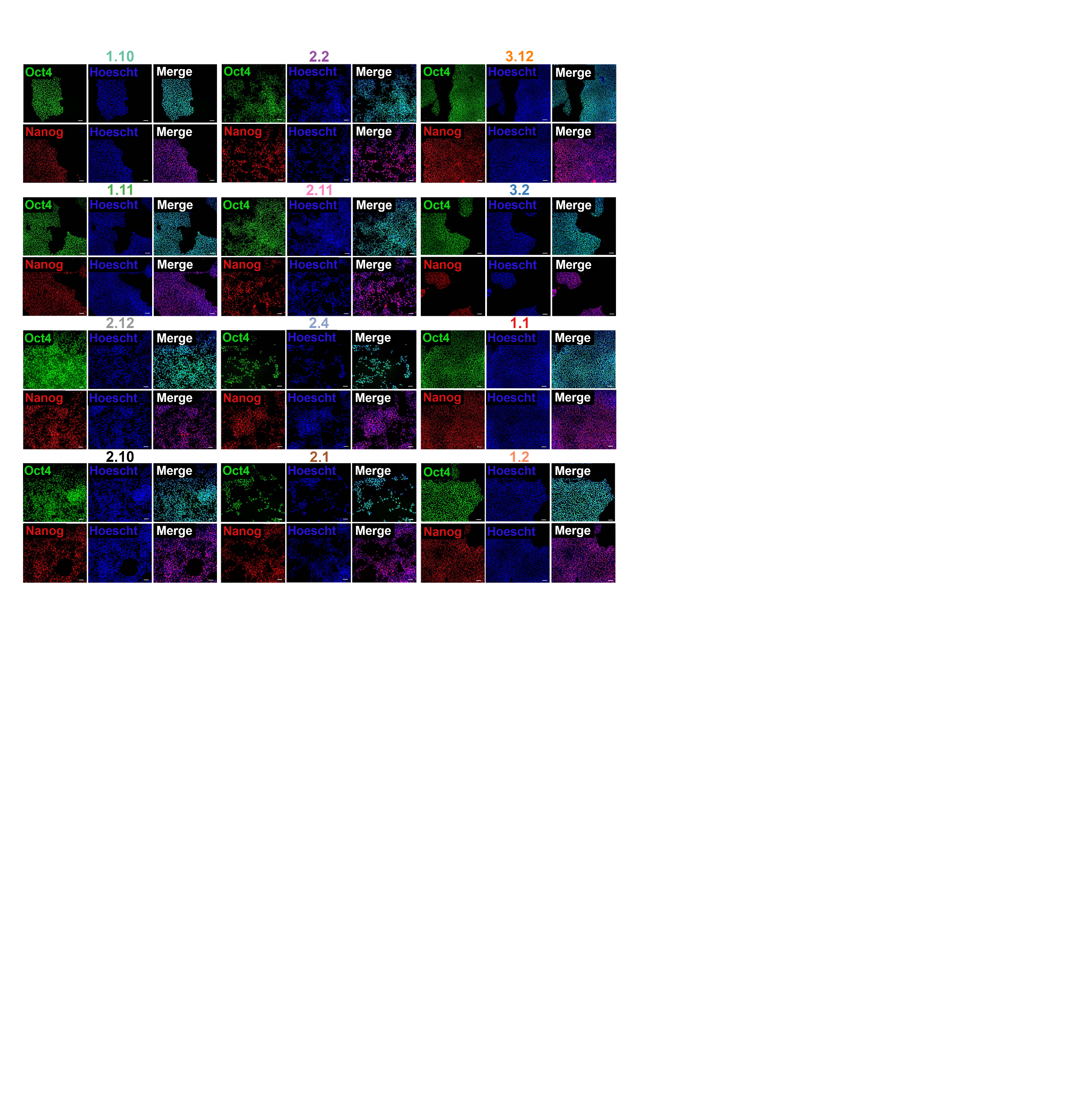

Figure S1 Selected clones iPSC heterozygous for p.T487I retain pluripotency markers

Twelve selected clones labelled for pluripotency markers Oct4 and Nanog. Scale bar is 100 microns.

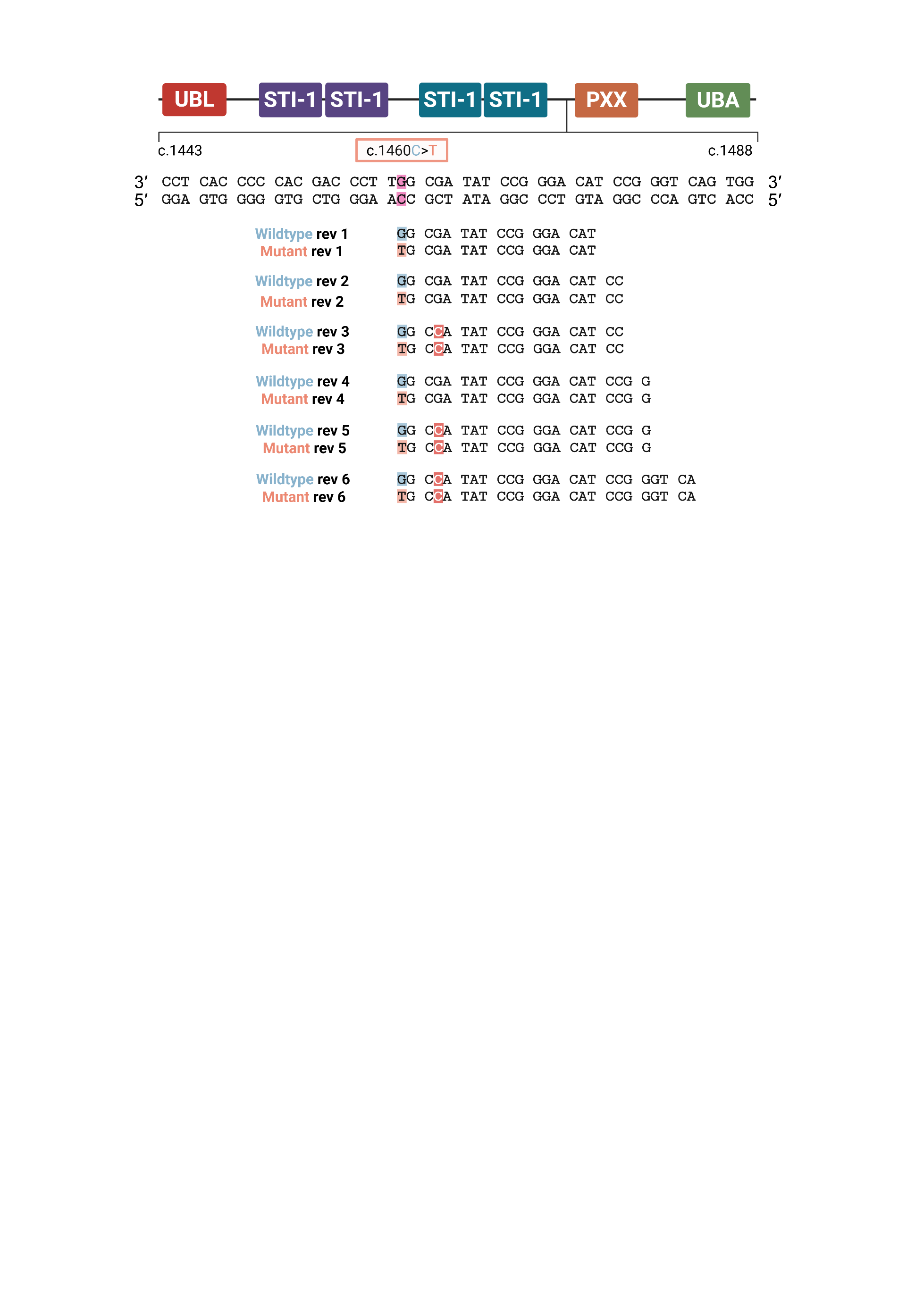

Figure S2 Primers targeting either the mutant *UBQLN2* p.T487I or wildtype allele in an allele-selective manner in the reverse (sense-binding) direction.

Allele-selective reverse primers were designed such that their 3' termini bind the p.T487I site (pink), creating a terminal mismatch when the primer binds the off-target allele. Allele-selective reverse primers differ in length, and whether or not they carry an introduced C:C mismatch (red) 3 base pairs from the 3' terminus. Allele-selective reverse primers were then paired with allele-agnostic primers predicted to share a similar annealing temperature. Schematic made in Biorender.

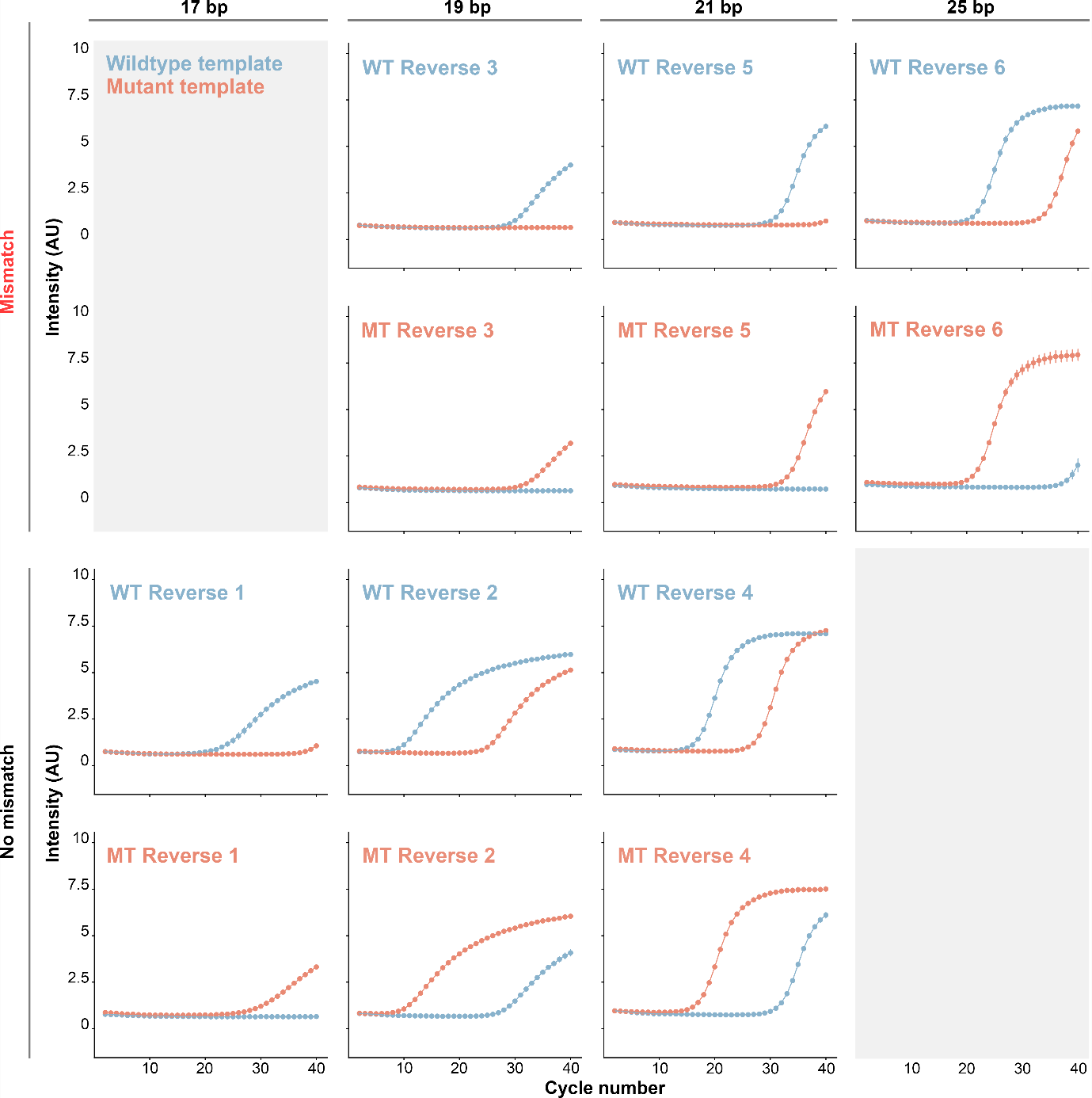

Figure S3 Amplification qPCR curve plots using wildtype- (WT) and p.T487I- (MT) selective primers, against wildtype and p.T487I *UBQLN2* plasmid DNA template.

Wildtype- and p.T487I-allele-selective reverse primers against mutant and wildtype *UBQLN2* plasmid DNA templates in a qPCR reaction. Mutant template in orange, Wildtype template in blue. Reactions contained 1068 pg of plasmid DNA at a primer annealing temperature of 65 °C. Data are mean +/- SEM from two technical repeats. Wildtype Reverse 4 and Mutant Reverse 4 were selected as hit primers (with their respective allele-agnostic primer partner) to generate a standard curve.

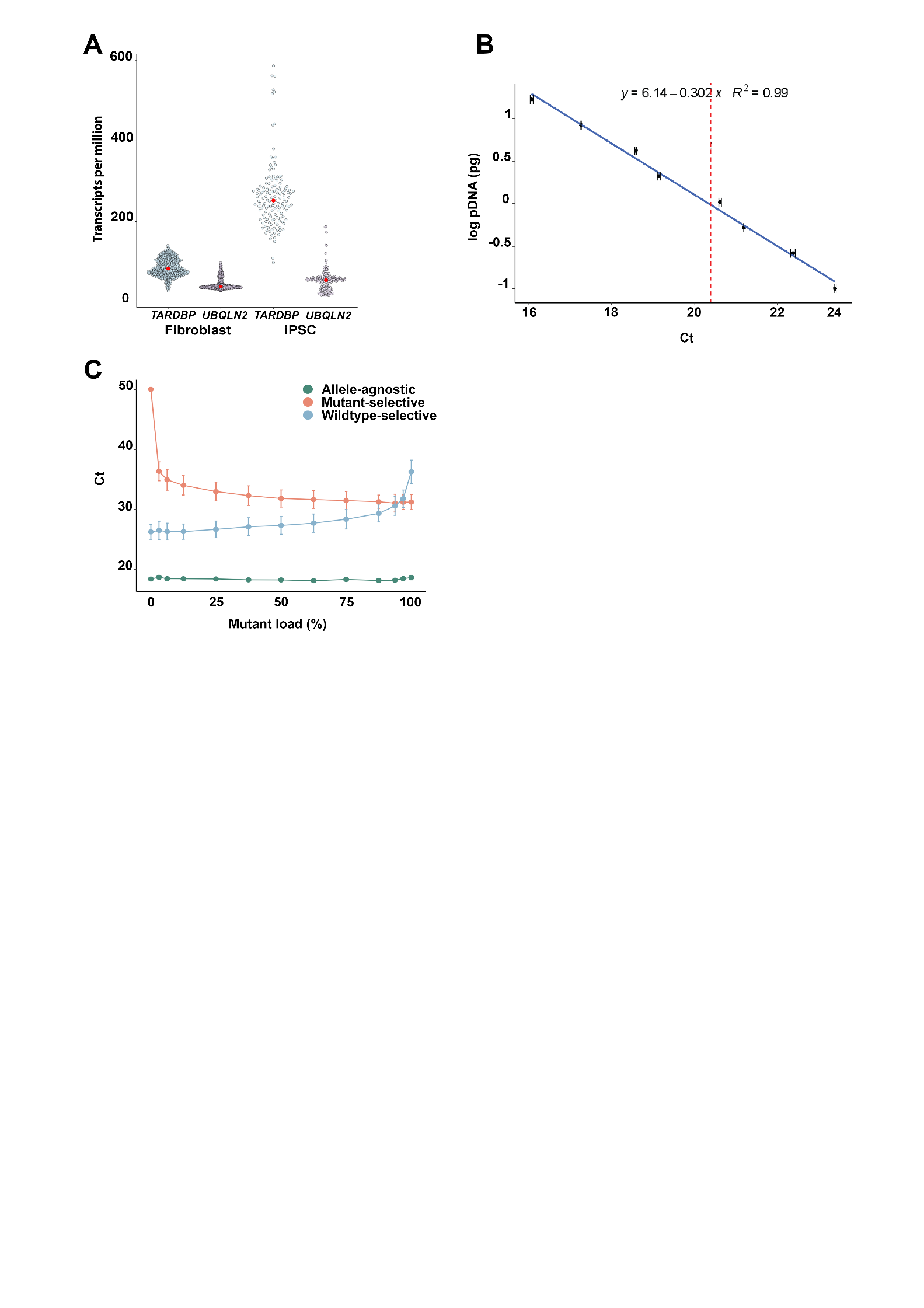

Figure S4 Development of the AS-qPCR assay

(A) Abundance of *UBQLN2* and *TARDBP* transcripts in HDF and HDF-derived iPSCs. HDF datapoints represent 504 entries from healthy HDFs deposited in GTEx, iPSC datapoints represent 145 entries from healthy HDF-derived iPSCs deposited in Stemformatics across 8 experiments. White point denotes mean of each group. (B) Ct values from allele-agnostic primer targeting *UBQLN2* by serial dilution of pDNA. Vertical red line demarcates Ct value for 10 ng of wildtype HDF cDNA. (C) Ct values for allele-agnostic, Mutant Reverse 4 and Wildtype Reverse 4 primer pairs across increasing *UBQLN2* p.T487I pDNA load. Data are mean +/- SEM from two independent replicates.
